# Impact of Long Term Integrated Nutrient Management Practices on Soil Bacterial Diversity: A 16S rRNA-Based Metagenomic Approach

**DOI:** 10.1101/2025.02.19.639006

**Authors:** Vishal Ahlawat, Navjeet Ahalawat, Nisha Boora, R S Dadarwal, Deepika Dhanda, Pooja Sangwan, Vipin Kumar, Rajiv Kumar, Parmod Kumar Yadav

## Abstract

The growing world’s population, expected to surpass 9 billion by 2050, demands a 70–100% increase in farm productivity, posing significant challenges for sustainable agriculture. Conventional farming, reliant on chemical fertilizers and pesticides, negatively impacts environmental sustainability and food quality. Metagenomics, a culture-independent technique, has revolutionized the study of microbial communities by enabling comprehensive analysis of microbial diversity and function. This study evaluated the effects of combined nutrient management practices—including farmyard manure (FYM), wheat straw, green manure, and NPK fertilizers—on soil bacterial diversity using 16S rRNA-based metagenomic analysis. Key findings revealed the dominance of bacterial phyla such as Proteobacteria, Actinobacteria, and Firmicutes, which play critical roles in nutrient cycling, organic matter decomposition, and plant growth promotion. Distinct taxonomic distributions were observed across the samples, with Proteobacteria and Firmicutes dominating in control (T1), NPK (T2), and NPK_Green Manure (T5) treatments, while a shift in phyla composition was noted in NPK_FYM (T3) and NPK_WheatStraw (T4). Genus-level analysis revealed that Pseudomonas and Bacillus species were abundant in control (T1) and NPK (T2), while Sphingomonas species were dominant in NPK_FYM (T3). Further diversity analyses revealed significant variation across the samples, with T3 showing the highest richness and control (T1) showing the lowest richness. The rarefaction curve confirmed sufficient sampling for capturing microbial diversity, while beta diversity analysis, including Principal Coordinate Analysis (PCA) and Bray-Curtis clustering, indicated distinct microbial community structures across different treatments. The combined use of organic amendments with NPK fertilizer enhanced microbial diversity and population, indicating their potential to support resilient and productive soil ecosystems.

## 1. Introduction

Soil is a dynamic and essential living system, critical for sustaining agricultural productivity, supporting plant and animal well-being, and preserving or improving water and air quality. It is widely acknowledged that various agricultural management practices, including crop rotation, tillage, crop residue management, irrigation and fertilization, have a profound influence on the physico-chemical characteristics of soil (Singh et al., 2011; Aziz et al., 2013). Soil fertilization is a time-honored agricultural practice for enhancing soil fertility, optimizing crop production, and boosting yields. In recent years, scientists have emphasized on studying the effect of various fertilization methods on soil microbes. The primary objective of soil fertilization systems is to provide essential nutrients to the soil that may be deficient or lacking, to make assure that plants have access to the essential elements for their healthy growth and maximum productivity (Jat et al., 2013). These systems encompass various practices, including the application of organic or inorganic fertilizers, the utilization of amendments to improve soil structure and nutrient availability, the implementation of precision techniques for targeted nutrient application, and the adoption of sustainable methods to minimize environmental impact (Cuartero et al., 2021).

Conventional soil fertility management practices typically rely heavily on chemical fertilizers rather than organic sources. The widespread use of these fertilizers can hinder nutrient absorption, degrade quality of soil, and lead to bigger issues such as eutrophication and greenhouse gas emissions (Zhu et al. 2016; Hartmann et al. 2014). In light of environmental challenges posed by fertilizers and the push for sustainable agriculture, research scholars are currently exploring the benefits of replacing fertilizers with organic sources of nutrients or integrating both organic materials and fertilizers to enhance nutrient levels while fostering a sustainable microbial ecosystem (Bhattacharyya et al. 2008; Babalola et al. 2009).

Soil microbes play a vital function in the ecosystem and act as soil health indicator. Recent advancements in molecular analysis have significantly enhanced our knowledge of plant-microbes interactions and their functions. Soil microbes are essential for maintaining soil fertility, as they actively contribute to key biological processes like nutrient cycling and organic matter decomposition. These processes are crucial for the efficient functioning of agricultural systems (Nicolas et al., 2012; Chaparro et al., 2012). Microbial-mediated processes are vital for carbon and nitrogen cycling, which are essential for the overall sustainability of agri-food systems (Govaerts et al., 2009; Osler and Sommerkorn, 2007). Historically, microbial community studies have consistently indicated that soil fertilized with organic manure exhibits greater bacterial and microbial diversity, regardless of experimental sites or climatic conditions. This increase in diversity is attributed to the nutrient enrichment provided by the organic sources of nutrients (Hamm et al. 2016; Chávez-Romero et al. 2016). In a comparative study, Chaudhry et al. (2012) observed that compost application significantly boosted the populations of certain bacterial phyla more effectively than chemical fertilizers, due to enhancements in soil carbon and nitrogen levels. While fertilization generally promotes microbial biodiversity, it is important to note that plants also play a critical role in shaping soil microbial communities by the mechanisms such as root exudation (rhizodeposition), as well as temperature and moisture regulation (Denef et al. 2009; Chen et al., 2023).

Numerous research have examined how agricultural practices affect the size, composition, and functionality of microbial communities in various crops, soils, and climates (Kihara et al. 2012; Lupwayi et al. 2012; Wang et al. 2012). Regardless of the techniques used, these results consistently indicate that cultivation practices influence soil microbial communities. Most research has focused on changes in composition of microbial communities and the fluctuations in the relative abundance of higher taxonomic groups, such as phyla or classes (Alsaedi et al., 2022). However, there is limited understanding of how long-term nutrient management practices impact lower ranking taxa, specifically at the genus and species level. These taxas are crucial for defining microbial communities (Hartmann et al., 2014; Jiménez-Bueno et al., 2016; Sun et al., 2021).

In this study, we sought to improve the analysis of microbial communities by identifying key taxa that influence their structure and correlating these taxa with variations in nutrient management practices. Our goal was to identify the key taxa affected by different agricultural methods and to leverage taxonomic, phylogenetic, and operational taxonomic unit (OTU) data to gain insights into how these factors impact the overall community structure.

## 2. Material and methods

### Sampling site description

Soil samples were collected from a long-term field experiment that commenced in Kharif 1985, located at the research farm of the Department of Agronomy at CCS Haryana Agricultural University (CCSHAU) in Hisar, India (29°08’58” N, 75°40’51” E). The climate of the area is classified as Bsh (Hot semi-arid) according to the Köppen Climate Classification System (Kottek et al., 2006), with an annual rainfall of 443 mm. Approximately 80% of this rainfall occurs between July and September, while the remainder falls from December to February. Summers are characterized by extreme heat, with maximum temperatures reaching up to 47°C, whereas winters are notably cold, with minimum temperatures dropping to 1°C. The soil in this region is classified as Typic Ustochrept, sandy loam soil. The long-term experiment employs a randomized complete block design featuring 12 treatments, each replicated four times. For the purpose of metagenomics study, five of these treatments have been selected. Detailed descriptions of the treatments can be found in Table 1.1. Each main plot measures 10 m by 8 m and includes various nutrient management treatments.

**Table 1.**
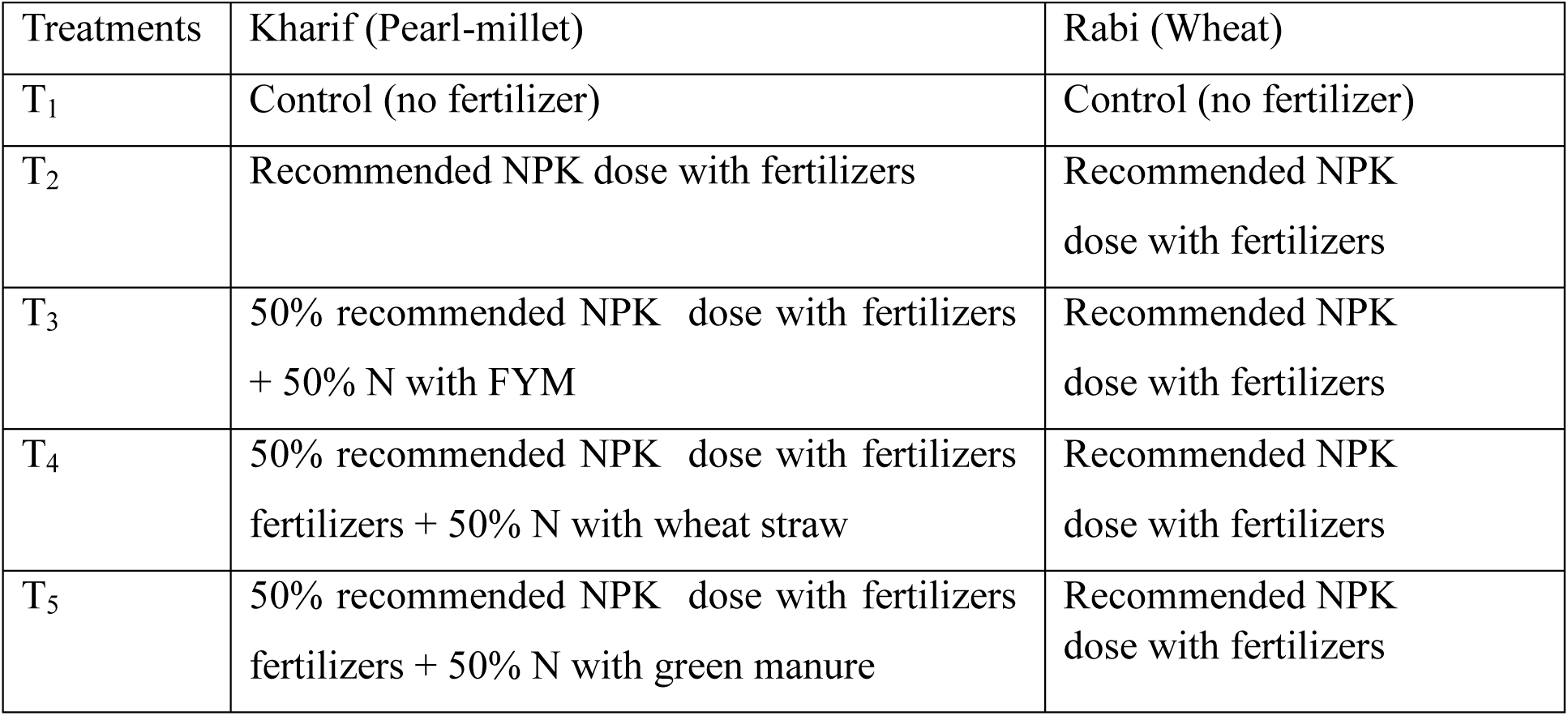
Details of the treatments.

Every year organic sources of nutrients including FYM, wheat straw and green manure were incorporated in the soil about 35-40 days before the sowing of pearl millet. Quantity of organic materials used to substitute the 50% of N was calculated on the basis N content of the material. The wheat straw was cut into small pieces and then incorporated into the soil. The dhaincha (Sesbania spp.) as green manuring crop was grown in a separate field and after about 45 days, it was harvested and chopped into small pieces and incorporated into the soil. Inorganic sources of nutrients were applied through Urea (46% N) and SSP (16% P_2_O_5_) fertilizers. The recommended dose of N in pearl millet crop was 125.0 kg ha^-1^ and that of phosphorus was 62.5 kg P_2_O_5_ ha^-1^. The total quantity of phosphorus and half of the nitrogen dosage, in accordance with the treatments, were applied at the time of planting, while the remaining nitrogen was applied following the first irrigation. For wheat crop the recommended dose of N and P was 150 kg ha^-1^ and 60 kg P_2_O_5_ ha^-1^ respectively. In pearl millet cultivation, the entire phosphorus amount and half of the nitrogen dose were applied at sowing, while the remaining nitrogen was added after the first irrigation.

The cropping pattern is an annual sequence of pearl-millet (***Pennisetum glaucum***) in kharif season and wheat (***Triticum aestivum* L**) in rabi season. Pearl millet (*Pennisetum glaucum*) is grown as a summer crop, typically planted in June and harvested in October. Following this, wheat (*Triticum aestivum* L.) is cultivated as a winter crop, which is planted from late October to early November and harvested in April. Soil samples were collected from a depth of 0-15 cm after the wheat harvest during the Rabi season of 2022-23.

### Sequencing and Metagenomic Sequence Analysis

#### DNA Extraction from Soil Samples

Using the MP Biomedicals FastDNA™ Spin Kit, genomic DNA was extracted from soil samples by the manufacturer’s instructions and kept at −80 °C. After isolation, Nanodrop and Agarose gel electrophoresis were used to assess the purity of the DNA, and a Qubit 4 Fluorometer (Thermo Fisher Scientific, USA) was used to confirm the DNA concentrations.

#### PCR Amplification and Sequencing

The V3-V4 region of the 16S rRNA gene was amplified via PCR, utilizing the extracted DNA as a template. Gene-specific primers 806R (GGACTACNNGGGTATCTAAT) and 341F (CCTAYGGGRBGCASCAG) were used for this process. The PCR conditions included an initial step at 98°C for 1 minute, followed by 30 cycles of 98°C for 10 seconds, 57°C for 30 seconds, and 72°C for 1 minute, concluding with a final extension at 72°C for 5 minutes. The quality of the purified PCR amplicons was assessed using a Bioanalyzer 2100. Subsequently, libraries were prepared with the TruSeq DNA PCR-Free Library Preparation Kit (Illumina). A Qubit 4 Fluorometer and a QuantStudio 5 Real-Time PCR system were employed to quantify the libraries. The qualified libraries were sequenced with a read length of 250 bp using paired-end chemistry on the Illumina NovaSeq 6000 platform.

#### Metagenomic Sequence Analysis

The sequence data underwent quality filtering in several stages. Initially, a “trim back valley filter” was used for 3’ trimming. Sequences were then sorted by barcodes, and chimeric sequences were identified using UCHIME. A feature table was generated to summarize the sequences linked to each sample. Representative sequences for each operational taxonomic unit (OTU) were selected for annotation, and a pre-trained naive classifier was used to classify each read or amplicon sequence variant (ASV). Phylogenetic relationships were inferred by aligning the representative OTU sequences with MAFFT, masking uninformative sites before creating the phylogenetic tree. The normalized data was utilized for further analyses of alpha diversity indices—such as observed species, evenness, Shannon diversity, and Faith’s phylogenetic diversity—using QIIME2. Beta diversity analysis, including dissimilarity matrices like Unweighted UniFrac and Weighted UniFrac, assessed differences in microbial communities between samples. Cluster analysis followed principal component analysis (PCA), and the results, along with taxonomy and tree data, were exported for further analysis in R.

## Results

A total of 409,439 16S rRNA sequences from five soil samples (T1, T2, T3, T4, and T5) were analyzed using the QIIME pipeline. Table 2 provides detailed information about the samples, including their conditions, high-quality reads, and GC content. Using the Greengenes database as a reference, 137,140 OTUs were identified across all five samples.

**Table 2.** Statistical Data for Samples T1, T2, T3, T4, and T5.

| Sample | Condition | Raw reads | HQ reads | Q20 (%) | GC content (%) |
| --- | --- | --- | --- | --- | --- |
| T1 | Control | 15526 | 14504 | 97.77 | 53.80 |
| T2 | NPK | 26977 | 25230 | 97.82 | 54.98 |
| T3 | NPK_FYM | 138889 | 130290 | 98.02 | 57.19 |
| T4 | NPK_WheatStraw | 116982 | 111512 | 98.06 | 56.49 |
| T5 | NPK_GreenManure | 134533 | 127903 | 98.04 | 55.47 |

Through QIIME, Proteobacteria, Firmicutes, and Actinobacteria were identified as the most abundant phyla across the five samples (T1, T2, T3, T4, and T5). Proteobacteria, which includes a wide range of gram-negative bacteria, was the most dominant phylum, followed by Firmicutes and Actinobacteria. In addition to the dominant phyla, other less abundant phyla were also detected in all five samples. These phyla include Chloroflexi, Acidobacteria, and Bacteroidetes (Figure 1A). In addition to the OTU table heat map, a relative abundance graph (Figure 1C) was generated to visualize the overall distribution of different phyla in the five samples. The analysis reaffirmed Proteobacteria as the dominant phylum across all samples. Notably, in samples T1, T2, and T5, Proteobacteria and Firmicutes constituted more than 60% of the total phyla. However, in samples T3 and T4, Proteobacteria and Actinobacteria were more abundant than Firmicutes, indicating a shift in the relative composition of phyla in these samples.

**Figure1:**
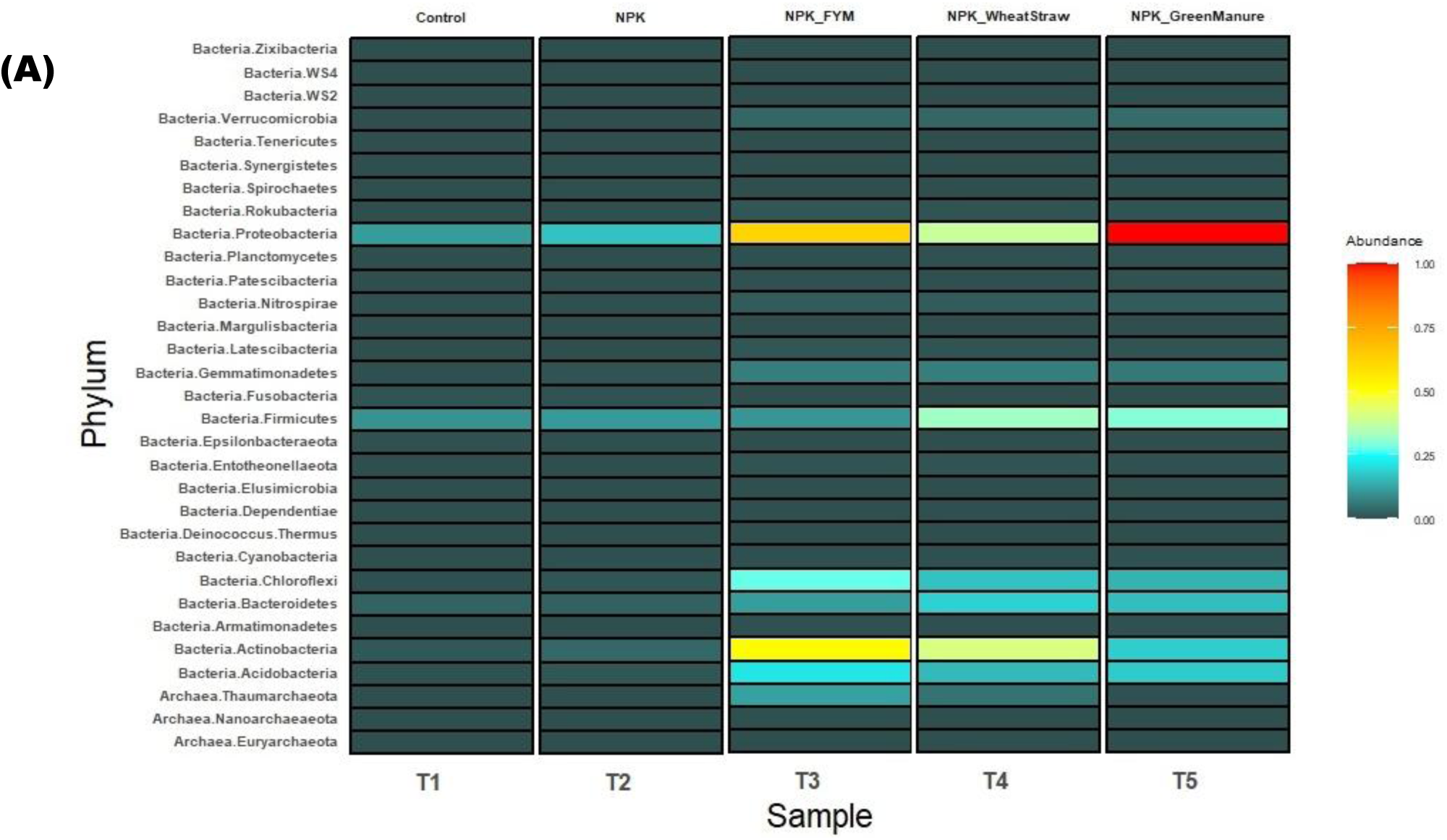
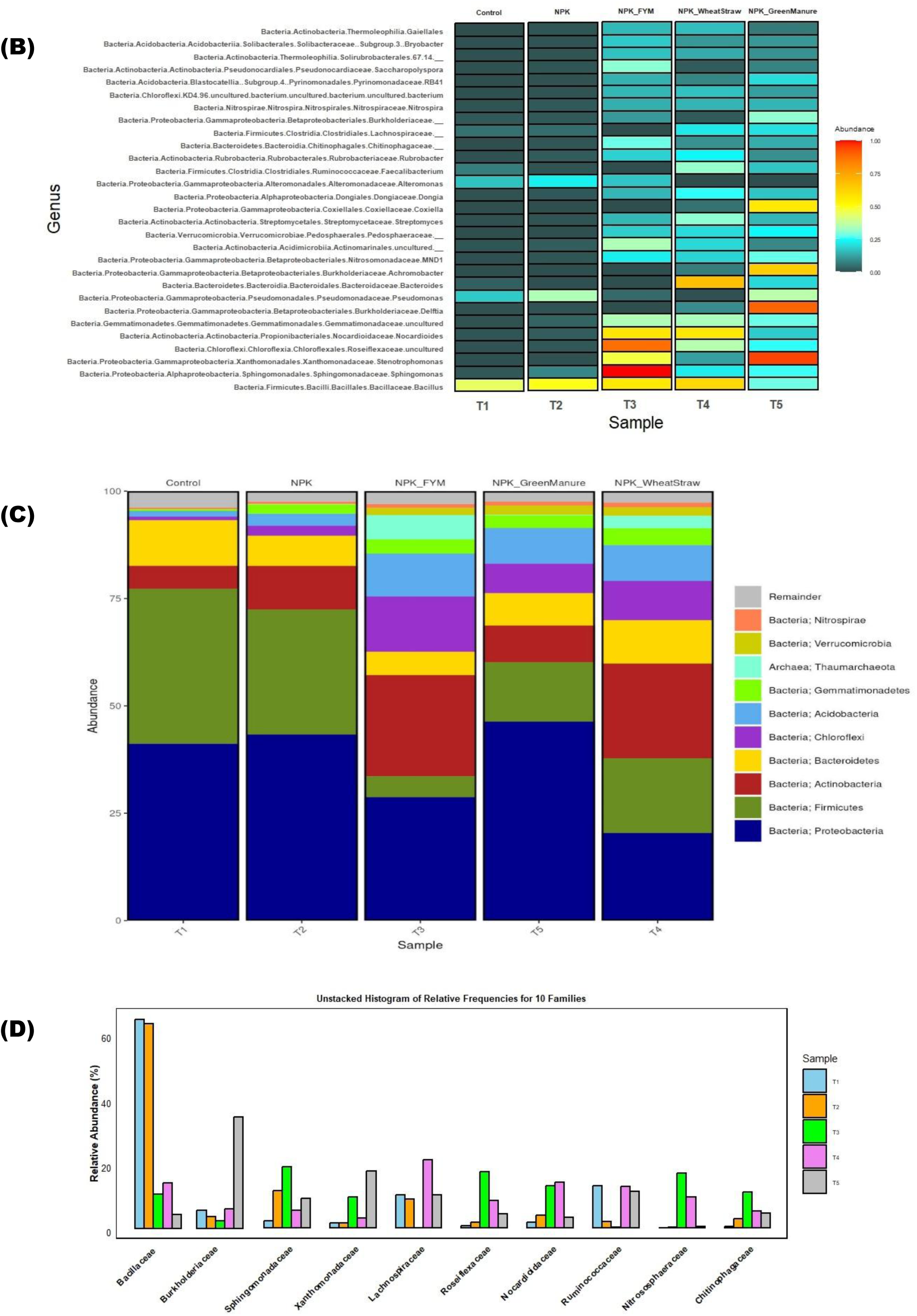
Microbial community composition of soil for five samples T1(Control), T2(NPK), T3(NPK_FYM), T4(NPK_WheatStraw) and T5(NPK_GreenManure) at **(A)**Phylum level and, **(B)** genus level. The heatmap illustrates the raw OTU counts for each sample, with colors representing the contribution of each OTU to the total OTU count in that sample. Blue indicates a low contribution of OTUs, while red signifies a high contribution. (C) displays the relative abundances of all phyla present in the soil samples. (Microorganisms with a relative abundance below 0.10% were classified as “Remainder.”). **(D)** Unstacked Histogram of relative frequencies for top 10 families among samples.

**Figure2:**
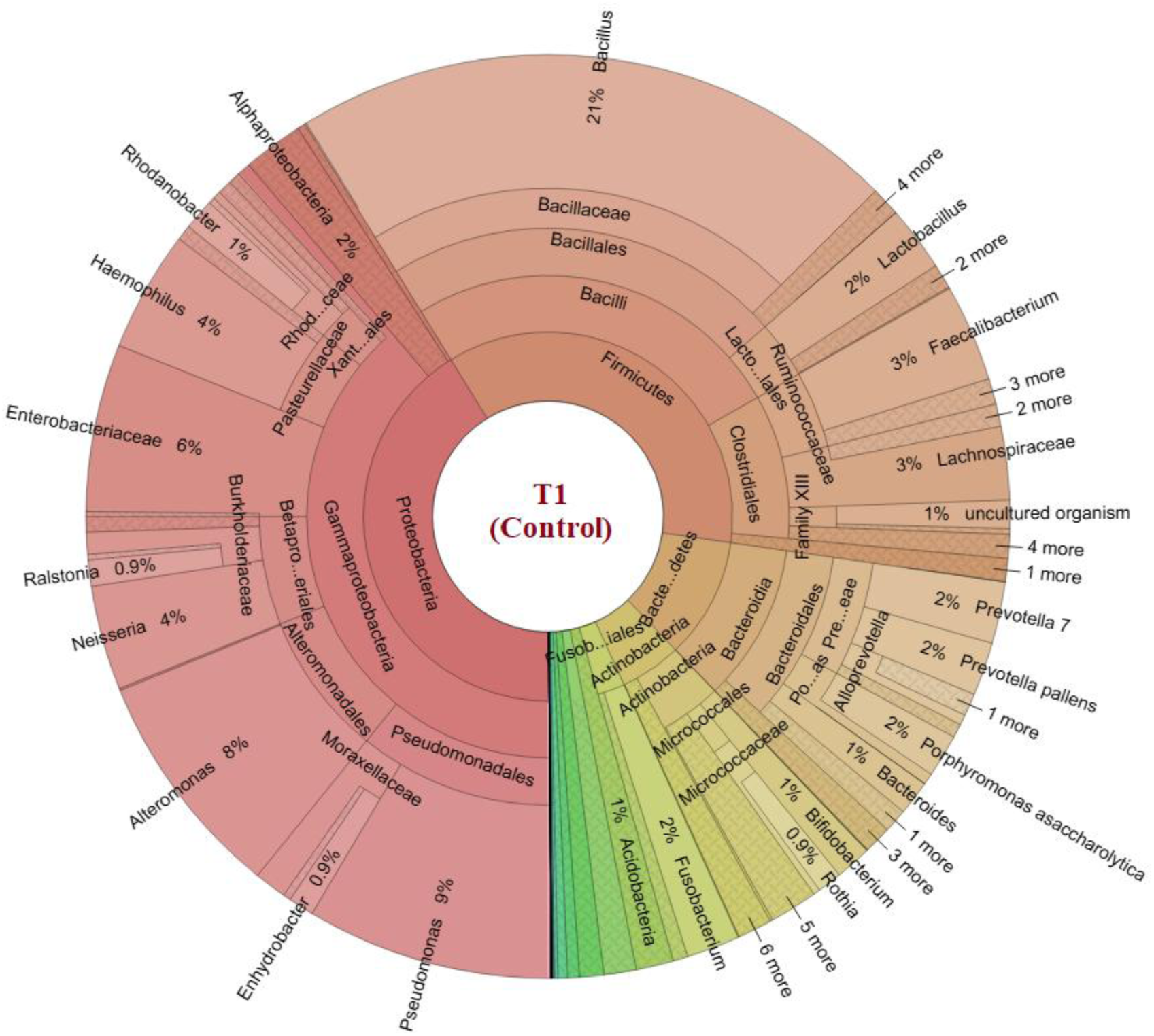
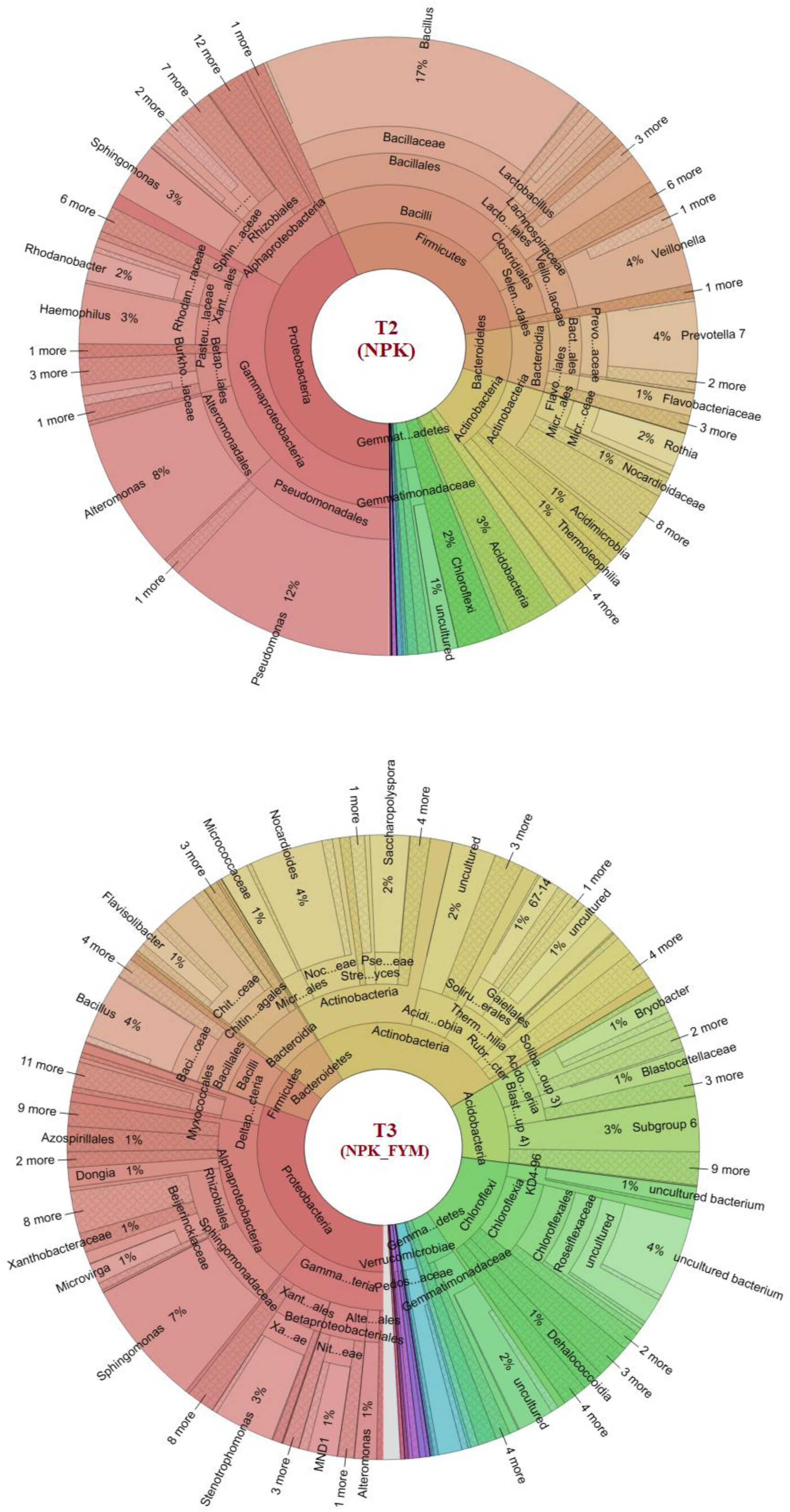
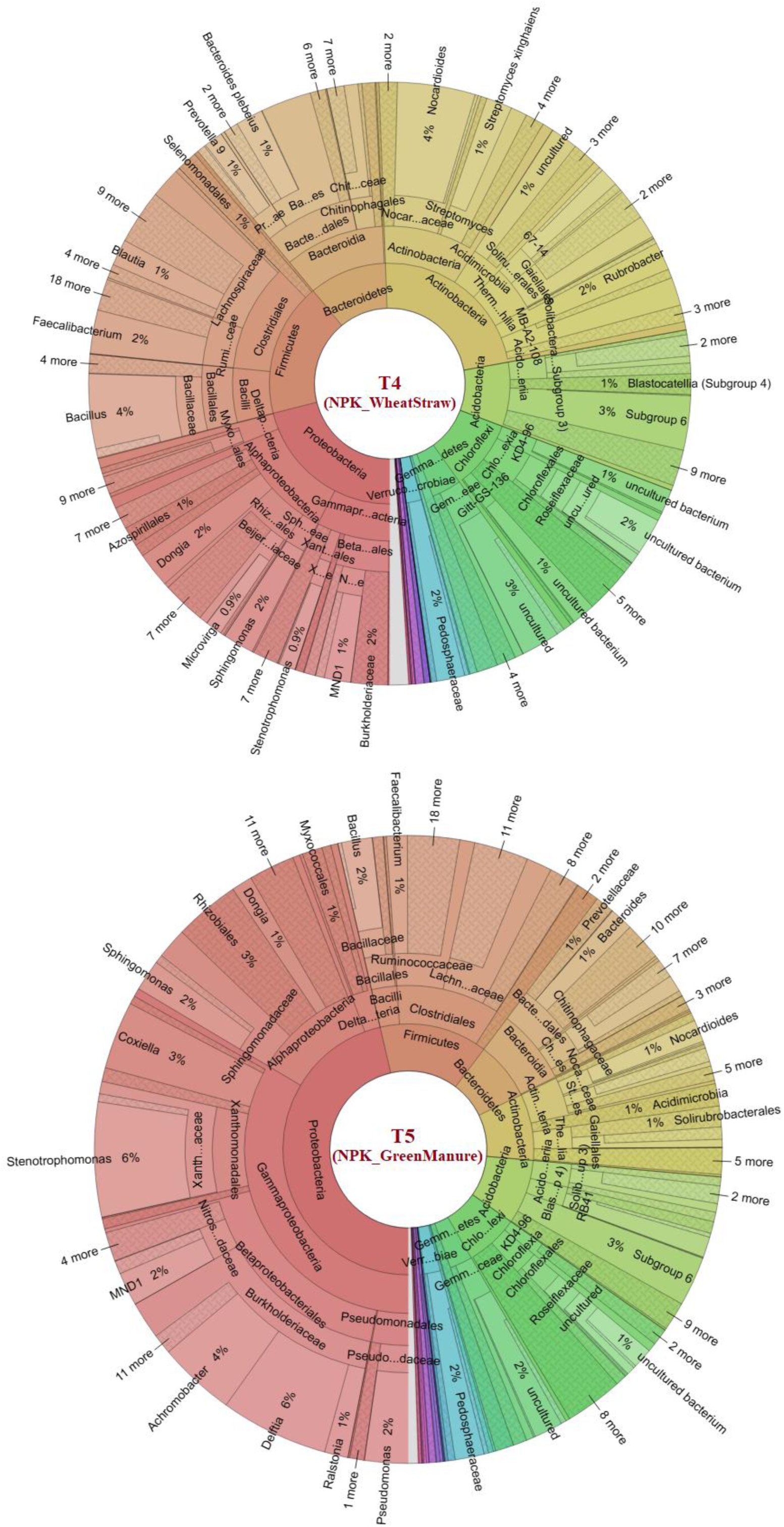
Krona graph displaying the taxonomy assignment at the class level for all samples.

Among the classes of Proteobacteria, Gammaproteobacteria exhibited the highest relative abundance, ranging from 7% to 38%. Sample T1 showed a higher relative abundance of Gammaproteobacteria, while T3 and T4 had higher levels of Alphaproteobacteria (56% and 54%, respectively) compared to the other treatments. The top 10 families with an average relative abundance greater than 2% were analyzed (Figure 1D). Samples T1 and T2 resulted in a notable increase in the relative abundance of Bacillaceae, while significantly reducing the relative abundances of Xanthomonadaceae, Roseiflexaceae, Nitrososphaeraceae, and Chitinophagaceae. Similarly, the relative abundances of Lachnospiraceae and Ruminococcaceae were significantly reduced in the T3 sample. In contrast, T5 showed a marked increase in the abundances of Bacillaceae and Xanthomonadaceae (Figure 1D).

In samples T1 and T2, the Proteobacteria were primarily classified as Gammaproteobacteria, specifically belonging to the order Pseudomonadales, family Pseudomonadaceae, and genus Pseudomonas. This taxonomic assignment suggests the presence of Bacillus species within the Firmicutes phylum in samples T1, T2, T3 and T4. In sample T3, the Proteobacteria were predominantly identified as Alphaproteobacteria, specifically associated with the order Sphingomonadales, family Sphingomonadaceae, and genus Sphingomonas. This finding indicates the prevalence of Sphingomonas species within the Proteobacteria phylum in sample T3. Besides Bacillus, sample T4 have significance abundance of Bacteroides and Nocardioides. On the other hand, in sample T5, the Proteobacteria were classified as Gammaproteobacteria, specifically assigned to the genus Stenotrophomonas, Delftia, Achromobacter and Coxiella are highly abundant (Figure 1B).

As a powerful visualization tool, KRONA presents a clear and concise representation of the species annotation analysis results. The visualization utilizes circles to depict different taxonomic ranks, arranged from the innermost to the outermost circle. Within each circle, the size of the sectors reflects the proportion of OTU annotation results for the specific taxonomic rank (Figure2). Notably, the phylum Proteobacteria appears to be enriched and dominant in samples T1, T2, and T5, collectively accounting for approximately 40% of all phyla. Conversely, the phylum Actinobacteria is relatively less abundant, comprising less than 10% of the total phyla in these samples. In contrast, samples T3 and T4 exhibit a comparable presence of Proteobacteria and Actinobacteria, with Actinobacteria representing 29% and 23% of all phyla, respectively. This suggests a more balanced distribution between these two phyla in these particular samples. Furthermore, the phylum Firmicutes exhibits varying abundance levels across the different samples. In T1 and T2 samples, Firmicutes constitute approximately 36% and 29% of all phyla, respectively. However, in the T5 sample, Firmicutes account for only 14% of all phyla, while Actinobacteria, Bacteroidetes, Acidobacteria, and Chloroflexi are found to be equally represented. Overall, the visualization provided by KRONA assists in comprehending and analyzing the species annotation analysis results by offering a visually intuitive representation of taxonomic ranks and their corresponding proportions in each sample.

### Alpha diversity

Alpha diversity is a measure of biodiversity that assesses the variety of organisms within a single ecosystem. The observed OTUs represent the richness, which counts the number of distinct taxa observed in a sample at a specific taxonomic level, typically at the species level. This provides a snapshot of the diversity present in each sample. On the other hand, the Shannon index is a more comprehensive estimator that takes into account both species richness (the number of different species) and evenness (the distribution of species within a sample).

The analysis revealed significant variations in alpha diversity among the five samples, highlighting how the total number of observed species ranged from 250 to 1250. Sample T3 had the highest number of observed species, almost reaching 1250, while sample T1 had the lowest count with only 260 species. Significant variations were observed in the alpha diversity indices among the five samples, with T1 and T2 displaying the lowest alpha diversity measures which means lowest richness, while T3 exhibited the highest alpha diversity measures, denotes highest richness (Figure 3A. A similar pattern was observed for the Shannon index (Figure 3B), where T1 shows the lowest diversity and evenness but here T3 and T4 both displayed higher diversity, indicating more evenly distributed species. Faith’s PD measures phylogenetic diversity, incorporating evolutionary relationships. T1 has the lowest phylogenetic diversity, while T3 and T4 are the most diverse (Figure 3C). Overall, these results conclude that T1 consistently shows the lowest diversity across all metrics, indicating it is less complex or diverse. T3 and T4 have higher richness, evenness, and phylogenetic diversity, suggesting they are more diverse and complex microbial communities. T2 and T5 fall in between, with moderate diversity. This suggests significant variability in diversity across the samples, which could relate to differences in experimental conditions.

**Figure3:**
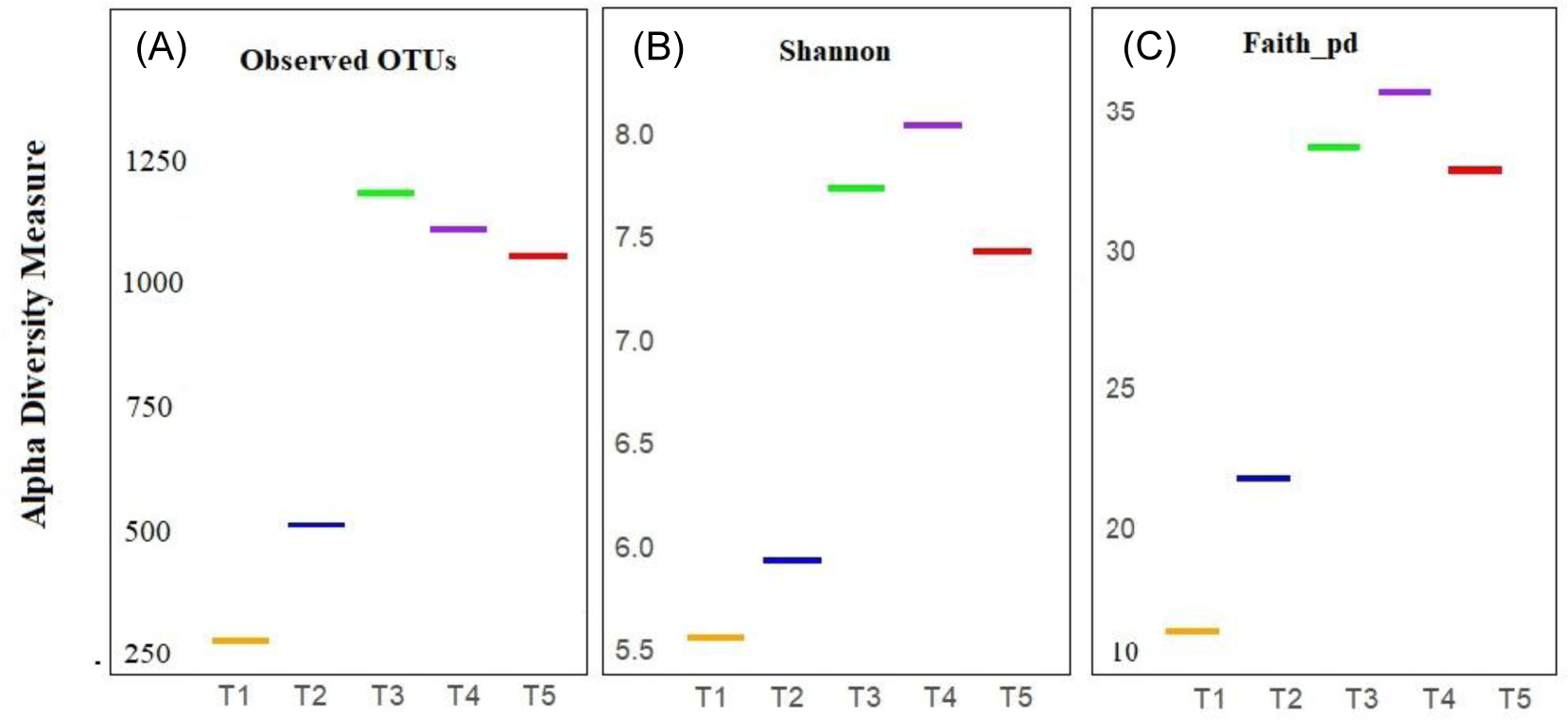
Alpha diversity measures for five samples (T1 to T5) across three metrics: (A) Observed OTUs, (B) Shannon index, and (C) Faith’s PD.

The rarefaction curve assists in assessing whether further sampling is necessary to obtain a more complete understanding of the species or taxa found within a particular population. It plots the number of distinct species or taxa (on the y-axis) against the number of sequences examined (on the x-axis). Continued sampling of the population will initially result in a sharp increase in the rarefaction curve. This reflects the detection of the most common and abundant species or taxa. The pronounced incline indicates the likelihood of uncovering many more species with further sampling. As more samples are collected, the rarefaction curve will begin to level off. This indicates that the sampling is increasingly uncovering the rarer species or taxa within the population. The curve will plateau once a significant fraction of the overall diversity has been sampled, leaving only the most uncommon species yet to be discovered. Figure 4 shows a rapid initial increase in the graph, followed by a gradual flattening of the curve. This pattern suggests that the most abundant and common microbial taxa have been well-captured through the sampling effort. However, there may still be some rare taxa that have not yet been detected.

**Figure4:**
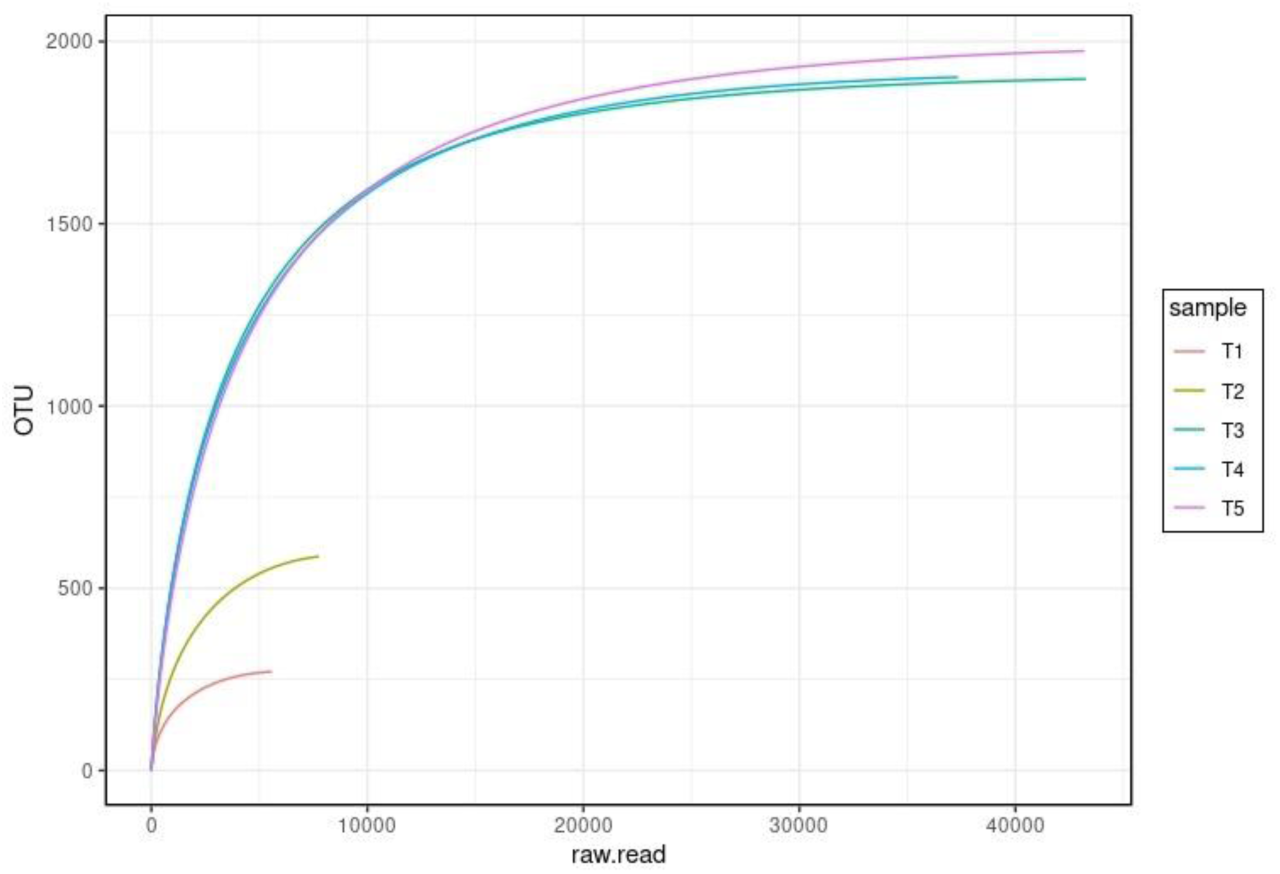
Rarefaction curve analysis of the five soil samples.

### Beta Diversity Analysis

The Beta diversity metric does not generate a value for each individual sample. Instead, it computes the distance between pairs of samples. The Bray–Curtis distance is a non-metric measure that quantifies compositional dissimilarity between two samples, taking into account the abundance data for each genus. This method is particularly suited for analyzing community data because it emphasizes differences in relative abundances rather than absolute values. QIIME utilizes Principal Component Analysis (PCA) plots to assess the distances between each sample. The closer the composition of the community among the samples, the nearer their corresponding data points will be on the PCA graph. The outcome of the principal component analysis is depicted in Figure 5A, where each point represents one of the samples. The results of the graphical representations on the PC1 (47.60%) and PC2 (26.11%) axes demonstrate that, sample T3 (NPK_FYM), T4 (NPK_WheatStraw) and T5 (NPK_GreenManure) stands apart from the other samples, while the T1 (Control) - T2 (NPK) and samples are located in close proximity to each other.

**Figure5:**
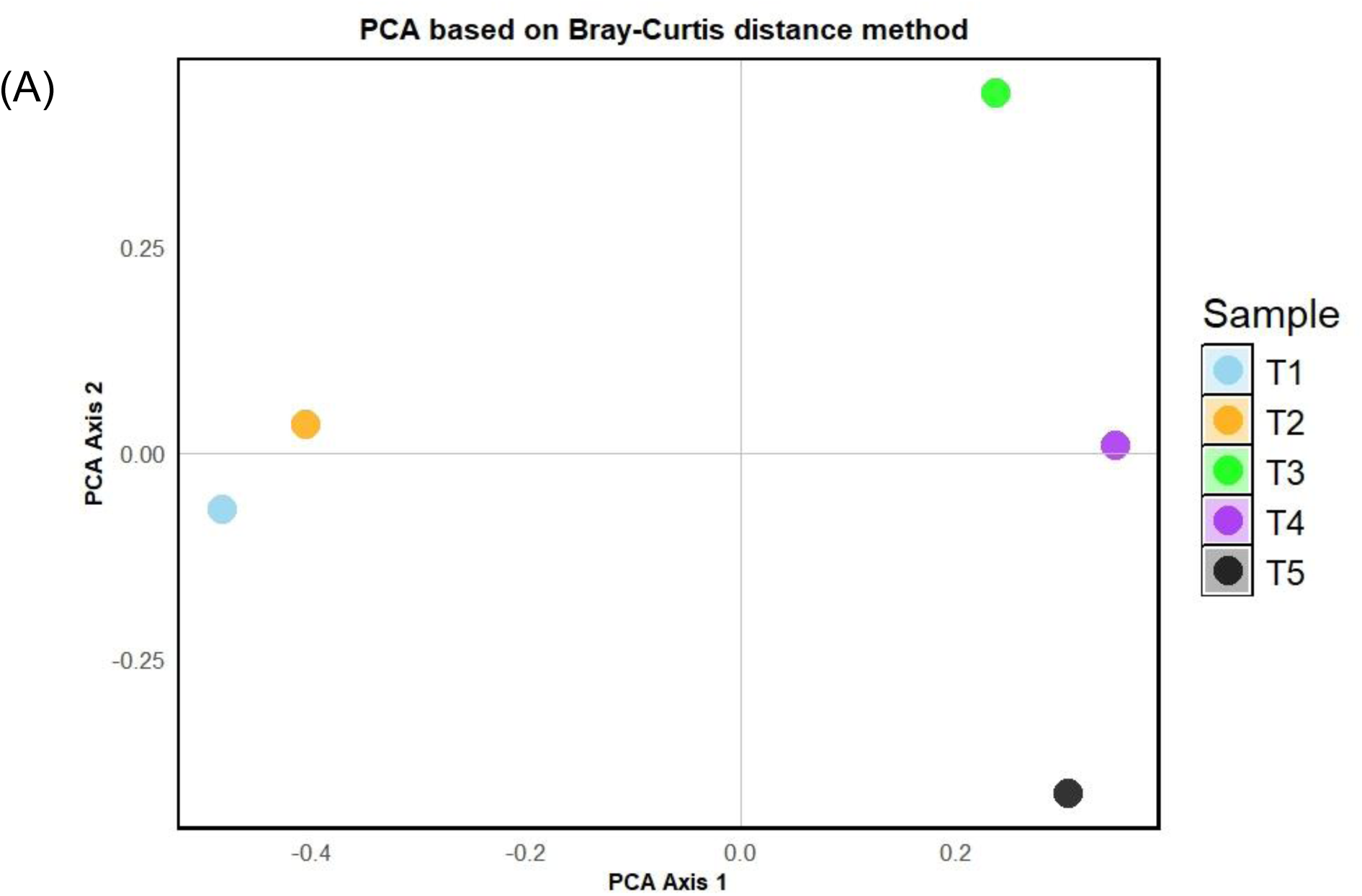
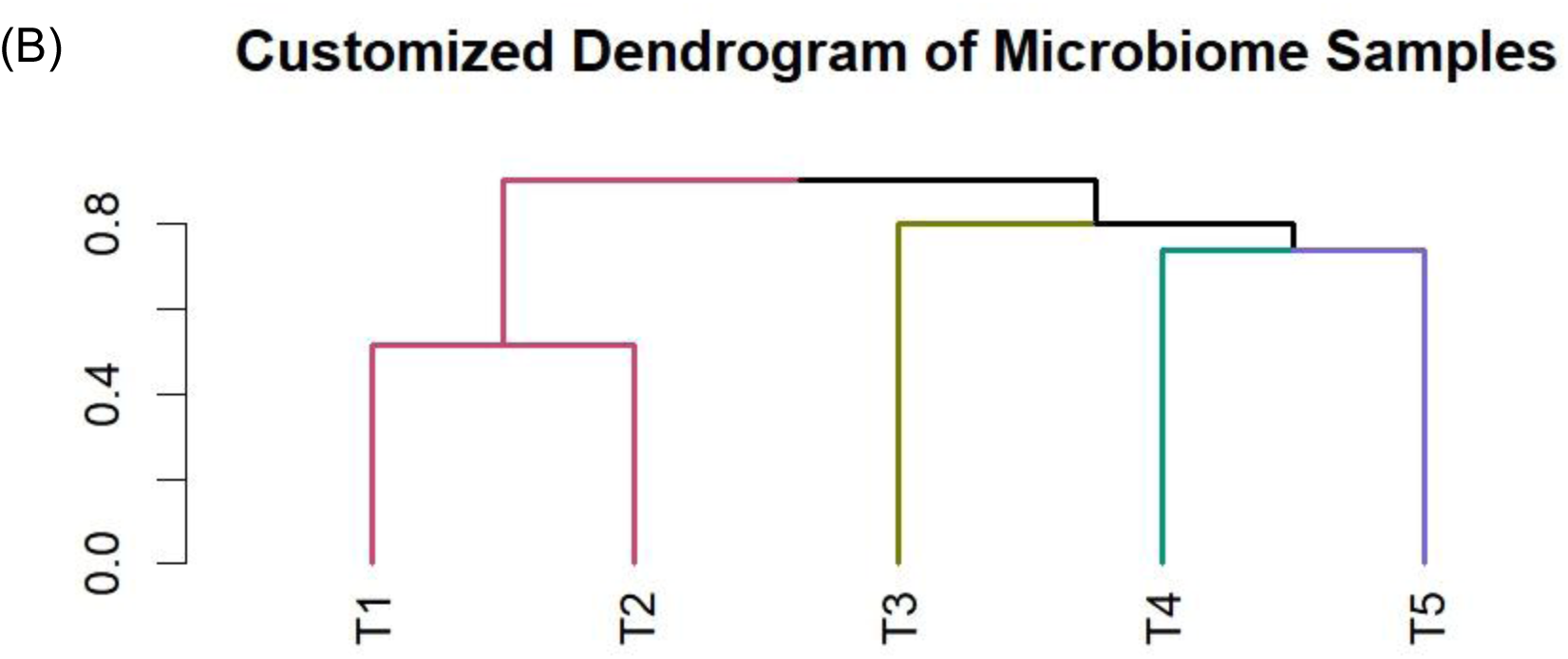
**(A)** Principal coordinate analysis (PCoA) was conducted using Bray–Curtis distances to assess the microbiota composition across different groups at the phylum level. **(B)** Cluster analysis was also performed based on Bray–Curtis distances at the genus level.

Cluster analysis based on Bray–Curtis distances at the genus level is a widely used method in ecological and microbial community studies to assess similarities or differences between samples. In this approach, abundance data of microbial genera across different samples are used to calculate pairwise Bray–Curtis distances. The resulting distance matrix is then subjected to clustering algorithms, such as hierarchical clustering or k-means clustering, to group samples based on their compositional similarities. Hierarchical clustering involves Ward’s method to construct dendrograms, providing a visual representation of sample relationships (Figure 5B). Overall observation suggests that the bacterial community structure was influenced in different sample types, indicating distinct differences between the bacterial community structures in the T1,T2 soil samples and the T3, T4 and T5 soil samples.

## Discussion

Microorganisms are everywhere in the biosphere, supporting life on Earth. Host-associated microorganisms, shaped by millions of years of co-evolution, are vital for their hosts’ fitness and health. Recent studies have underscored the significant influence of plant-associated microbes, collectively known as the phyto-microbiome, on plant health and productivity (Pattnaik et al., 2021). This has given rise to various strategies aimed at leveraging the phyto-microbiome to enhance resource availability, improve resource use efficiency, and provide protection against pests and pathogens, thereby supporting sustainable agriculture. However, traditional microbiological methods face limitations, as a majority of microorganisms remain unculturable (Li et al., 2017, Masuda et al., 2024,). Metagenomics now enables a comprehensive understanding of the structural and functional profiles of bacterial, archaeal, and fungal communities in both soil-based and soilless agricultural systems (Nwachukwu & Babalola 2022). This knowledge holds immense potential for advancing sustainable agricultural practices by leveraging microbial communities to optimize productivity and ecosystem health (Enebe & Babalola 2020, and Iquebal et al., 2022).

The utilization of organic amendments in agriculture has gained significant attention due to its potential to improve soil fertility, enhance nutrient availability, and promote sustainable agricultural practices (Sabale et al., 2019, Schmidt et al., 2019). Our research evaluated how the combined utilization of FYM (Farmyard Manure), wheat straw, green manure, with NPK fertilizer impacts the diversity and population of soil bacteria. We employed metagenomic analysis based on the 16S rRNA gene and our findings indicated variations in both the composition and abundance of soil bacteria in all the samples.

Our study found that Proteobacteria, Firmicutes, and Actinobacteria dominated all analyzed samples, in addition to other phyla such as Chloroflexi, Acidobacteria, and Bacteroidetes. Their prevalence suggests their adaptability to a wide range of agricultural management practices and their significant roles in nutrient cycling and organic matter decomposition. The presence of NPK fertilizer, either alone or with green manure, seemed to favour the growth and proliferation of Proteobacteria and Firmicutes. This observation aligns with previous studies indicating the resilience of these phyla to conventional agricultural practices. Interestingly, in the NPK with FYM and wheat straw sample, Proteobacteria and Actinobacteria exhibited high abundance. Bacterial species within the phylum Proteobacteria play a significant role in soil functions, including nutrient cycling and the maintenance of soil health. These gram-negative bacteria are among the most abundant in agricultural soils. Additionally, many Proteobacteria promote plant growth by producing growth-promoting substances and suppressing plant diseases. Their ability to degrade organic matter and pollutants plays a crucial role in nutrient cycling and maintaining soil health (Kumar *et al.,* 2021, Patil et al., 2020). Actinobacteria are gram-positive bacteria that play a role in the biogeochemical cycling of nutrients and produce a variety of secondary metabolites. They possess the capability to produce biologically active secondary metabolites, and it has been reported that approximately 16,500 compounds derived from Actinobacteria exhibit antibacterial properties against pathogenic bacteria. Actinobacteria also play a role in soil structure formation by producing hyphae that help create soil aggregates, improving water infiltration and aeration (Kundu 2018, Prabhu et al., 2017, Trivedi et al., 2022, Yurgel et al., 2018, Youssef & Elshahed 2009).

Firmicutes, another important phylum, contribute to nutrient cycling through the decomposition of organic matter. Certain Firmicutes exhibit antagonistic activities against plant pathogens, providing protection to plants and promoting their health. Furthermore, Firmicutes have demonstrated potential in bioremediation processes due to their ability to degrade a diverse array of organic pollutants (Patil et al., 2020, Zhou et at., 2023). Bacteroidetes are also highly abundant in soil and anaerobic environments. They inhabit the rhizosphere of plants, contributing to soil health, the degradation of organic polymers, and nutrient cycling. Their populations tend to increase with higher soil nutrient levels or fertilization (Badapanda *et al.,* 2017, Sahu et al., 2017, Oliveira et al., 2022).

Several bacteria were identified across five nutrient management treatments, illustrating the diverse and multifunctional nature of soil microbial communities. Several genera contribute significantly to plant growth promotion through nutrient solubilization, phytohormone production, and protection against pathogens. Notable among these are Tumebacillus, Bacillus, Pseudomonas, Sphingomonas, and Streptomyces, which also exhibit biocontrol properties (Oteino et al., 2015, Raimi et al., 2023). Nitrogen-fixing bacteria, such as Bradyrhizobium and the Allorhizobium-Neorhizobium-Pararhizobium-Rhizobium complex, play crucial roles in symbiotic nitrogen fixation with legumes, while Pseudomonas contributes through its nitrogen-fixing capabilities and other functions (Ahemad & Kibret, 2014, Rashid et al., 2016). Phosphate solubilizers like Bacillus, Pseudomonas, and Streptomyces enhance phosphorus availability, a key factor in energy transfer and root development. Decomposers such as Clostridium sensu stricto 1 and Solirubrobacter recycle nutrients and improve soil structure by breaking down organic matter. Other beneficial genera, including Microvirga, Kribbella, and Nitrospira, support plant health indirectly by maintaining microbial diversity. However, the presence of pathogenic genera like Nocardioides, Corynebacterium 1, and Ralstonia highlights the importance of balanced microbial populations to mitigate plant diseases. Genera with unknown or multifunctional roles, such as Nordella, Aeromicrobium, and Ammoniphilus, are less understood but are likely to have context-dependent ecological contributions. This variability underscores the complexity of soil ecosystems and their pivotal role in sustainable nutrient management (Saeed et al., 2021, Barbaccia et al., 2022, Wei, et al., 2024).

The genera Enhydrobacter and Moraxella are found to be unique under T1. Both genera are reported to have roles in environmental and nutrient cycling, but their specific roles in plant-microbe interactions remain less understood. They might act as generalists or have functions yet to be fully characterized. Research highlights that many bacteria with undefined functions may play roles in soil health and microbial diversity, contributing indirectly to plant productivity (Rodrigues et. al. 2024 and Hao et. al. 2024)

Under T2 treatment, Methylobacterium; known as a nitrogen fixer and carbon sequesterer, it promotes plant growth by enhancing nutrient uptake and contributing to nitrogen availability in the soil. It is also linked with stress resistance in crops. Acetobacter plays a key role in nitrogen fixation, particularly under low oxygen conditions, and is also involved in soil carbon dynamics. Studies demonstrate the ability of Methylobacterium to fix nitrogen in the phyllosphere and its potential to be used as a bioinoculant in crops. Similarly, Acetobacter has been studied for its nitrogen fixation potential in tropical and subtropical cropping systems (Rodrigues et. al. 2024 and Hao et. al. 2024, Reis et al., 2012). The role of Prevotella in plant systems is not well-defined, but related species are involved in the degradation of organic matter, suggesting potential roles in carbon cycling. Prevotella species are recognized for their microbial fermentation abilities in organic environments (Hao et. al. 2024, Trivedi et al., 2022, Yurgel et al., 2018).

In T3 treatment, as an unclassified genus, KCM-B-112 may play niche roles in microbial ecosystems. Its function is hypothesized to contribute to microbial diversity and resilience in soil ecosystems. Studies on microbial diversity emphasize the importance of even poorly understood taxa in maintaining ecosystem functions (Blake et al., 2021, Hao et. al. 2024, Zolti, et al., 2020).

The genera Megasphaera and Succinivibrio were unique to T4 treatment. These genera are known as decomposers, these genera assist in breaking down organic material, contributing to nutrient cycling in the soil. Ruminococcaceae UCG-014; functions remain unclear, but other members of the family are important for cellulose degradation and nutrient release. Research supports the roles of these genera in carbon cycling, specifically their contributions to organic matter decomposition in soil (Arrobas et. al. 2024, Fierer et al., 2007, Hao et. al. 2024, Sharuddin et al., 2022, Timofeeva et al., 2023).

The Hydrogenophaga, Thauera, Brevibacterium, Bacteroidetes bacterium OLB10, and Lentimicrobium genera were found exclusively in T5 treatment. Hydrogenophaga; a plant growth promoter, aiding in nutrient uptake and possibly influencing root architecture. Thauera; functions as a nitrogen fixer and contributes to nitrogen cycling in soils. Brevibacterium; a phosphate solubilizer, enhancing phosphorus availability to plants. Bacteroidetes bacterium OLB10; Its exact role is unclear, but related genera are known for organic matter decomposition. Lentimicrobium; Includes species beneficial for anaerobic digestion and organic matter transformation. These genera have been shown to contribute significantly to soil fertility and plant growth through nutrient solubilization and organic matter decomposition (Rodrigues et. al. 2024, Hao et. al. 2024, Jeyanthi et al., 2018, Martínez-Viveros et al., 2010, Vestergaard et al., 2017, Ye et al., 2024).

The combined application of farmyard manure (FYM), wheat straw, and green manure, along with NPK fertilizer, influenced the diversity and population of soil bacteria. The incorporation of organic amendments appeared to enhance bacterial diversity, highlighting the potential of organic amendments in promoting a more resilient and productive soil microbial community. These findings contribute to our understanding of the effects of combined nutrient management practices on soil bacterial communities and emphasize the importance of considering organic amendments for sustainable agriculture practices. Further studies are warranted to explore the responses of specific bacterial taxa and their functional roles in the context of combined nutrient management strategies.

## Conclusion

In conclusion, with the world’s population growing, we need new ways to farm better. As highlighted in this study, the interplay between microbial communities and nutrient management strategies underscores the potential for optimizing soil health and crop productivity. The investigation into the combined utilization of organic amendments and NPK fertilizer offers valuable insights into the intricate dynamics of bacterial populations within soil. Proteobacteria, Firmicutes, and Actinobacteria emerge as pivotal players in these microbial consortia, adapting and growing under varying treatment conditions. The integration of organic amendments such as FYM and wheat straw emerges as a promising avenue for enhancing bacterial diversity, potentially supporting the resilience and functionality of soil ecosystems. In a world where agricultural sustainability is of paramount importance, these findings contribute to the evolving blueprint for future farming practices, advocating for a balanced coexistence between human needs and the intricate microbial world beneath our feet.

